# Functional retinal imaging using adaptive optics swept-source OCT at 1.6MHz

**DOI:** 10.1101/420240

**Authors:** Mehdi Azimipour, Justin V. Migacz, Robert J. Zawadzki, John S. Werner, Ravi S. Jonnal

## Abstract

Objective optical assessment of photoreceptor function may permit earlier diagnosis of retinal disease than current methods such as perimetry, electrophysiology, and clinical imaging. In this work, we describe an adaptive optics (AO) optical coherence tomography (OCT) system designed to measure functional responses of single cones to visible stimuli. The OCT subsystem consisted of a raster-scanning Fourier-domain mode-locked laser that acquires A-scans at 1.64*MHz* with a center wavelength of 1063*nm*, and an AO subsystem providing diffraction-limited imaging. Analysis of serial volumetric images revealed phase changes of cone photoreceptors consistent with outer segment elongation and proportional to stimulus intensity, as well as other morphological changes in the outer segment and retinal pigment epithelium.

## 1 Introduction

Vision begins when photons are absorbed in the photoreceptor outer segment, initiating the biochemical process of phototransduction. In blinding retinal diseases such as retinitis pigmentosa and age-related macular degeneration, vision is lost when these cells become dysfunctional. Current methods for diagnosing and assessing retinal disease, such as examining the appearance of the retina in clinical images and assessing visual function with clinical exams, are effective after extensive pathological changes, but not in the earliest stages of disease. Adaptive optics (AO) flood imaging [1], conventional optical coherence tomography (OCT) [2, 3, 4], and full-field OCT [5] have revealed changes in the human photoreceptor outer segment (OS) in response to visible stimuli. Here, we describe an OCT imaging system that leverages the three-dimensional cellular resolution of AO and OCT[6, 7] and the speed of a Fourier-domain mode-locked (FDML) laser [8, 9]. This enabled us to resolve cone photoreceptors in three dimensions and characterize changes in single cones OS morphology evoked by impulse-like bleaching flashes. To our knowledge, the instrument described here is the fastest AO-OCT system to date.

## 2 Methods

### 2.1 AO-OCT System

A schematic of the AO-OCT system is shown in Fig. 1(A). The AO-OCT system consisted of two main parts: a swept source (SS)-OCT subsystem and an AO subsystem based on a Shack-Hartmann wavefront sensor (SHWS) and voice-coil based deformable mirror (DM). The swept-source was a Fourier-domain mode-locked (FDML) laser (FDM-1060-750-4B-APC, OptoRes GmbH, Munich, Germany) operating at an A-scan rate of 1.64*MHz* [8, 9]. A Michelson interferometer configuration with three 50:50 fiber couplers was used to balance spectra of two detection channels. This topology is advantageous in suppressing the relative noise intensity (RIN) from the laser [10]. The exposure level of the imaging beam was limited to 1.8*mW*, which is within the limit determined by the ANSI standard for the safe use of lasers, and the measured sensitivity of the system was –85*dB*. The sample arm of the system consisted of pairs of spherical mirror telescopes in an out-of-plane configuration (Fig. 1(B)) [11] to correct beam distortions and astigmatism that otherwise accumulate as light is relayed off-axis by multiple in-plane spherical mirrors. The scanning system contained a resonant scanner (SC-30, Electro-Optical Products Corp., Ridgewood, NY, USA) oscillating at 5*kHz* in the horizontal direction which allowed 160 A-scans in each B-scan, and a galvanometer scanner in the slower vertical direction. The scanner configuration, in concert with the A-scan rate, permitted acquisition of 32 volumes per second over a field of view of 1° × 1°. Table 1 summarizes the significant characteristics of the system and data acquisition settings during imaging. The axial resolution of the OCT system was estimated experimentally by placing a flat mirror in place of the subject’s eye and measuring the full width at half maximum (FWHM) of the PSF. As is shown in Fig. 2(B), the axial resolution was 10.8*µm* in air which corresponds to 7.8*µm* in tissue (n = 1.38). Sensitivity roll-off was determined by moving the reference arm while a flat mirror was placed in the sample arm. Based on the roll-off in Fig. 2(C), sensitivity was reduced by 6*dB* approximately 2*mm* from the zero path length.

The AO subsystem incorporated a SHWS consisting of a 20 × 20 lenslet array (Northrop-Grumman Corp, Arlington, VA,USA) in front of a sCMOS camera (Ace acA2040-180km; Basler AG), and a high-speed deformable mirror (DM-97-15; ALPAO SAS, Montbonnot-Saint-Martin, France). The wavefront beacon source was a 840*nm* superluminescent diode (Superlum Diodes Ltd, Cork, Ireland), with power measured at the cornea of 20*µW*. The system provided diffraction-limited imaging for a 6.75*mm* diameter beam by measuring and correcting ocular aberrations in closed-loop at a rate of 15*Hz*, yielding a theoretical lateral resolution of 3.2 *µm*. Custom software controlled the AO [12] and OCT data acquisition which were developed in Python and LABVIEW (National Instruments, Austin, Texas, US), respectively. Most of the OCT signal processing was done in MATLAB computing software (The MathWorks, Inc., Natick, MA, USA).

**Table 1:** Specifications of the AO-FDML system and scanning parameters during imaging.

|  |  |
| --- | --- |
| Laser center wavelength | $1063nm$ |
| Spectral bandwidth (FWHM) | $78nm$ |
| Laser A-scan rate | $1.64MHz$ |
| B-scan rate | $5kHz$ |
| Volume rate | $32Hz$ |
| Optical power at cornea | $1.8mW$ |
| Axial resolution in air | $10.8\mu m$ |
| Measured sensitivity | $-85.4dB$ |
| Laser phase noise (rms) | $2.6mrad$ |

**Figure 1:**
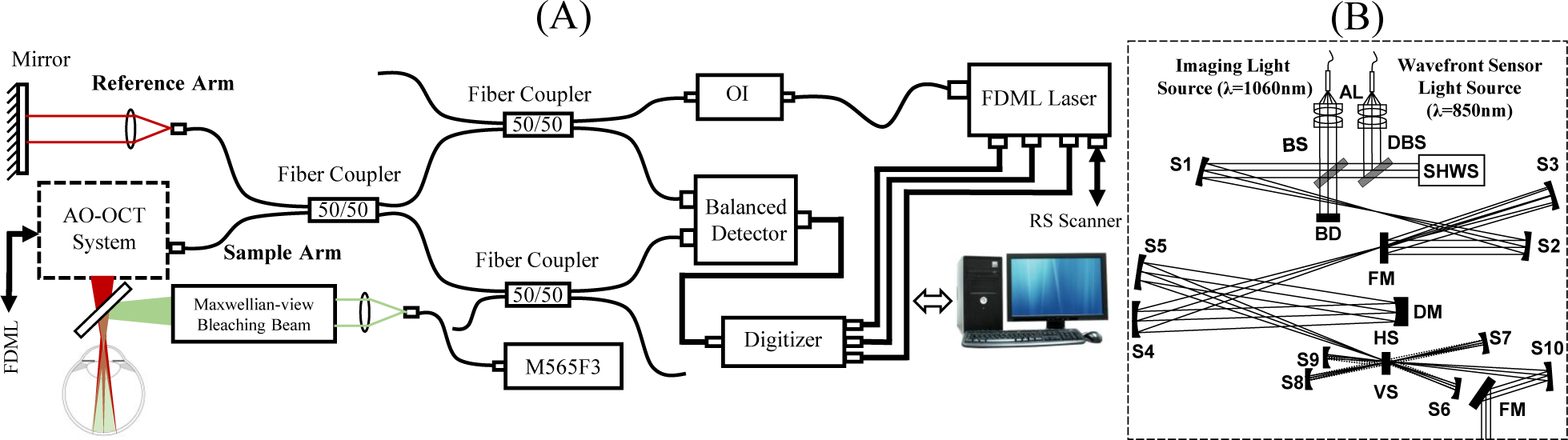
**(A)** Schematic of the AO-FDML OCT imaging system integrated with Maxwellian-view optical system for bleaching photoreceptors. **(B)** An expanded view of the AO scanning system: DM, deformable mirror; SHWS, Shack-Hartmann wavefront sensor; AL, achromatic lens; S, spherical mirror; FM, flat mirror; BS, beam splitter; DBS, dichroic beam splitter; HS, horizontal scanner; VS, vertical scanner; BD, beam dump; OI, optical isolator.

**Figure 2:**
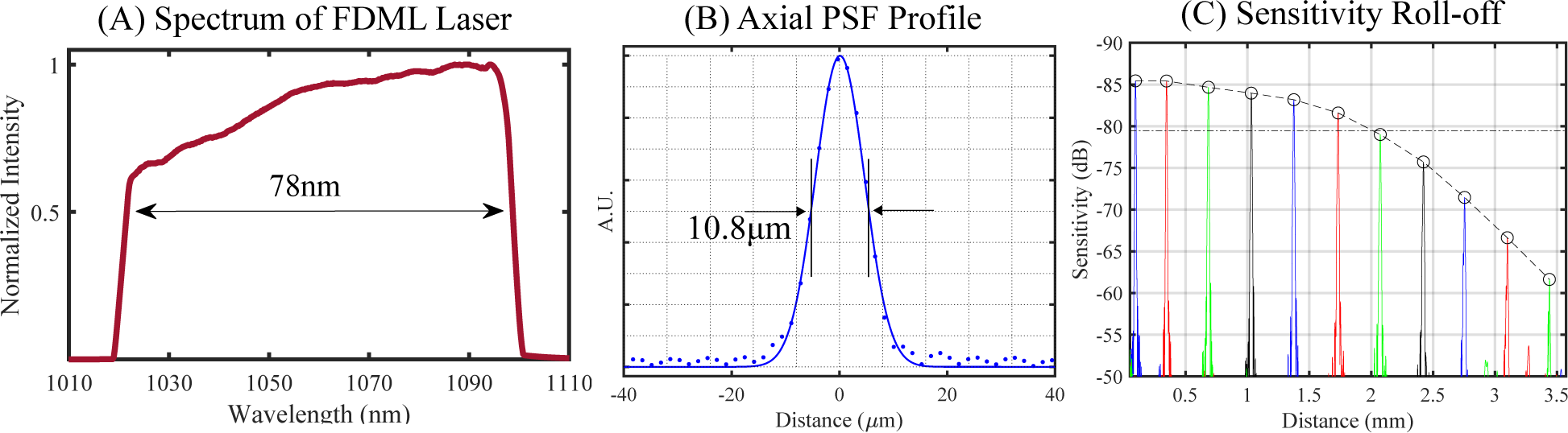
**(A)** Spectrum of FDML laser. **(B)** Axial point spread function and **(C)** sensitivity roll-off of the imaging system.

### 2.2 Imaging Protocole

Two subjects, free of known retinal disease, were imaged after obtaining informed consent. Each subject’s eye was dilated and cyclopleged by instilling topical drops of 2.5% phenylephrine and 1% tropicamide. All procedures were in accordance with the tenets of the Declaration of Helsinki and were approved by the University of California, Davis Institutional Review Board. To reduce head movement and stabilize pupil position during retinal imaging, a bite-bar and a forehead-rest were employed and assembled on a motorized X-Y-Z translation stage to precisely adjust the position of the subject’s pupil in the center of the imaging system entrance pupil. During imaging, a calibrated fixation target was employed to position the eye at specified retinal locations as well as to reduce eye movements. For functional imaging, subjects were dark-adapted for 15 minutes and then imaged for 10 seconds at a retinal location 2.5 degrees temporal to the fovea, where the expected cone row-spacing was ≈ 5.0*µm* [13]. At the 2-second baseline a 10*ms* visible flash was delivered. A bandpass filter centered at 555*nm* with 20*nm* bandwidth was placed in front of the bleaching light source, which was a fiber-coupled LED (M565F3, Thorlabs, NJ). This light spectrum provides almost identical bleaching efficiency for both L and M cones. Flash intensity was modulated in order to bleach between 1.8% and 70% of L/M photopigment.

### 2.3 Signal processing

A strip-based registration method [14, 15] was implemented to track individual cones in the volume series. First, the volumetric images were segmented axially and the inner-outer segment (IS/OS) and cone outer segment tips (COST) layers were automatically identified and projected. The *en face* projection of the cone mosaic from a single volume and an average of 30 motion-corrected volumes is shown in Fig. 3. For each series, a single IS/OS projection was selected as a reference image, and the remaining projections were divided into strips of between 5 – 11 pixels of height and registered to the reference. Cones were automatically identified in the reference image. Segmentation and lateral registration together permitted 3D tracking of single cones over time. Time-series of the complex axial signal (M-scans) of each cone were recorded. As the phase of the OCT signal provides a sub-resolution measure of the object’s motion[16], we recorded the phase difference between the reflection at IS/OS and COST[15], which provides an estimate of OS length change that is immune to artifacts of axial eye movement:

**Figure 3:**
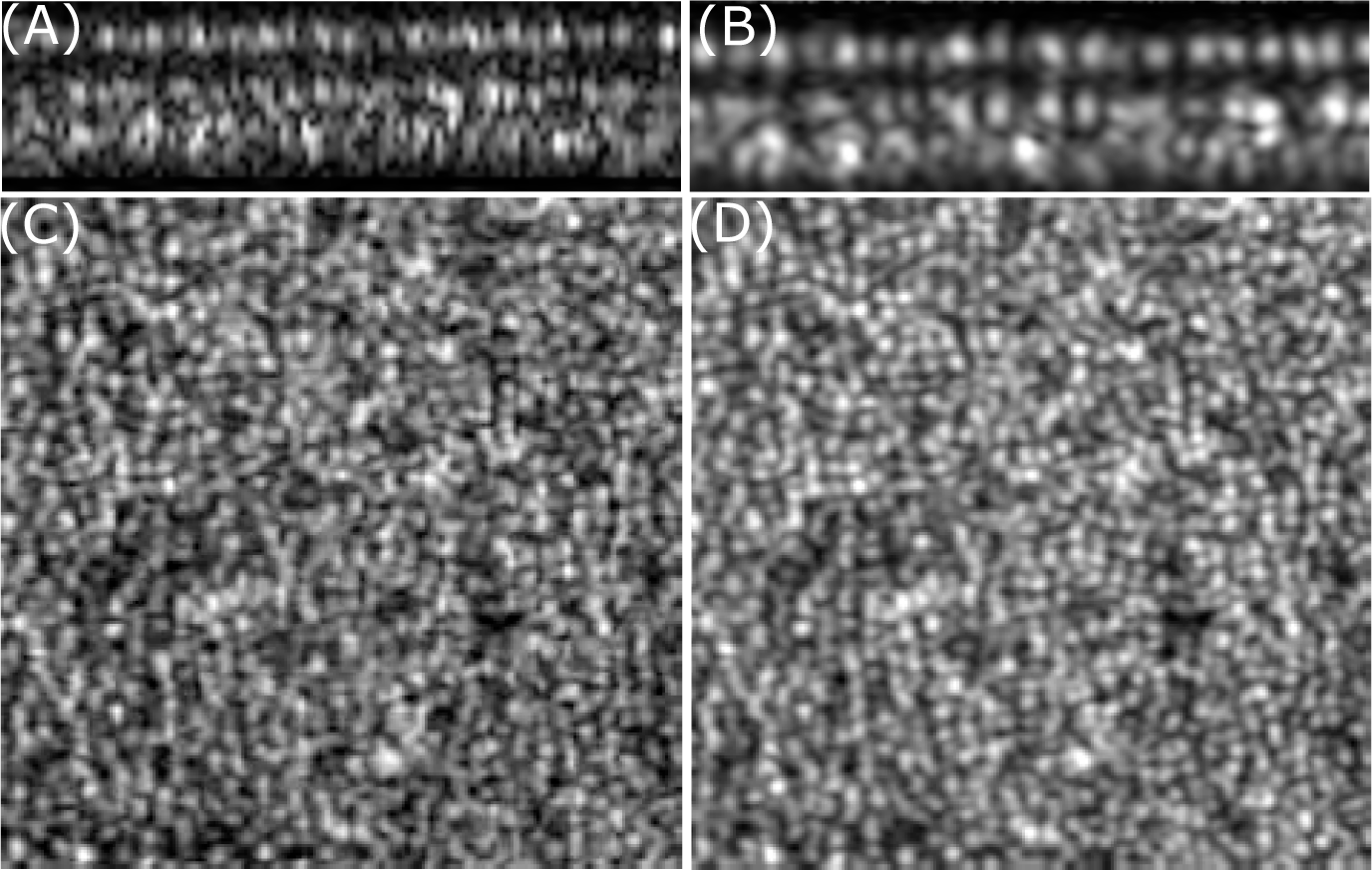
Strip-based registration permits averaging of AO-OCT volumes. Top panel shows **(A)** single B-scan and **(B)** average of 30 B-scans. En-face projection of cone mosaic from a **(C)** single and **(D)** average of 30 motion-corrected volumes of a 1° degree patch acquired at 2.5° temporal from the foveal center.

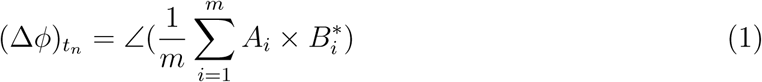

where *A* is a complex number corresponding to OCT signal measured at reflection from COST, *B*^*^ is the complex conjugate of OCT signal measured at reflection from IS/OS, and *m* is the number of *A*-scans recorded within each single cone. After subtracting the phase values from the initial phase difference, (Δ*ϕ*)_*t*_0__, and unwrapping them, the physical OS length change was measured using equation 2:

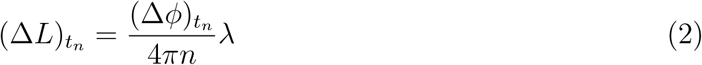

where *n* is the refractive index of the OS (∼ 1.38) and *λ* is the wavelength of the imaging light source (1063*nm*).

## 3 Results and Discussions

A response of a single cone to 70 percent photopigment bleaching stimuli is shown in Fig. 4(A). *En face projections* of the cone’s neighborhood are shown at the top, with the cone marked by a colored box indicating the time interval during which the cone was imaged. Next, the amplitude of the cone’s M-scan is shown, with the stimulus flash indicated by a green line. Changes in the cone’s axial profile subsequent to the flash are visible: the IS/OS reflectivity decreases; the distance between COST and IS/OS appears to increase; and structural changes are visible in other layers. At the bottom, the phase difference between IS/OS and COST is shown as a function of time. Responses of a single cone to three different flash intensities are shown in Fig. 4(B). Each response is the OS elongation corresponding to the measured phase difference between IS/OS and COST. It is apparent that the magnitude of elongation varies with stimulus strength, as does the time required for the cone to recover its baseline length. This observed elongation of foveal/parafoveal cones, and the dependence of elongation on stimulus intensity, is qualitatively consistent with previously reported elongation of peripheral cones using full-field OCT with computational aberration correction [5]. We hypothesize that the mechanism of this elongation is osmotic swelling of the outer segment, as it is consistent with observations made using conventional OCT images in human rods [4] and mouse rods, in which elongation is suppressed in mice lacking transducin [17]. In addition to OS elongation, changes in the axial morphology of cones were observed subsequent to the stimulus flash. Representative M-scans of single cones for photopigment bleaching percentages of 1.8, 7 and 70 are shown in Fig. 5. A common observation was the appearance and/or movement of an extra band between IS/OS and COST, indicated by red arrows in Fig. 5. The reflectivity of this band and its maximal axial distance from IS/OS appear to be proportional to the bleaching light intensity, and it is most evident in the 70% bleaching trials (5 C), where it moves half the length of the OS within 1-2 seconds. Generally, OCT signals are attributed to refractive index mismatch, and the movement of this band is consistent with the movement of a refractive index boundary. Such a boundary could be generated, for instance, by an abrupt change in disc spacing or concentration of a visual cycle intermediate. The observation is also consistent with coherent effects of modulation in disc spacing [17]. Another common observation was a change in the appearance of the space distal to COST, including the subretinal space (SRS) and RPE. The RPE band appears to split, with its apical portion moving inward, toward COST. If melanosomes are an important source of RPE scattering [18], the observed movement could be an indication of melanosome movement into the apical part of the RPE cell. This is consistent with light-driven translocation of melanin observed in amphibians [19], but not previously reported in mammals. It may also be a consequence of inward water movement across the Bruchs-RPE complex.

**Figure 4:**
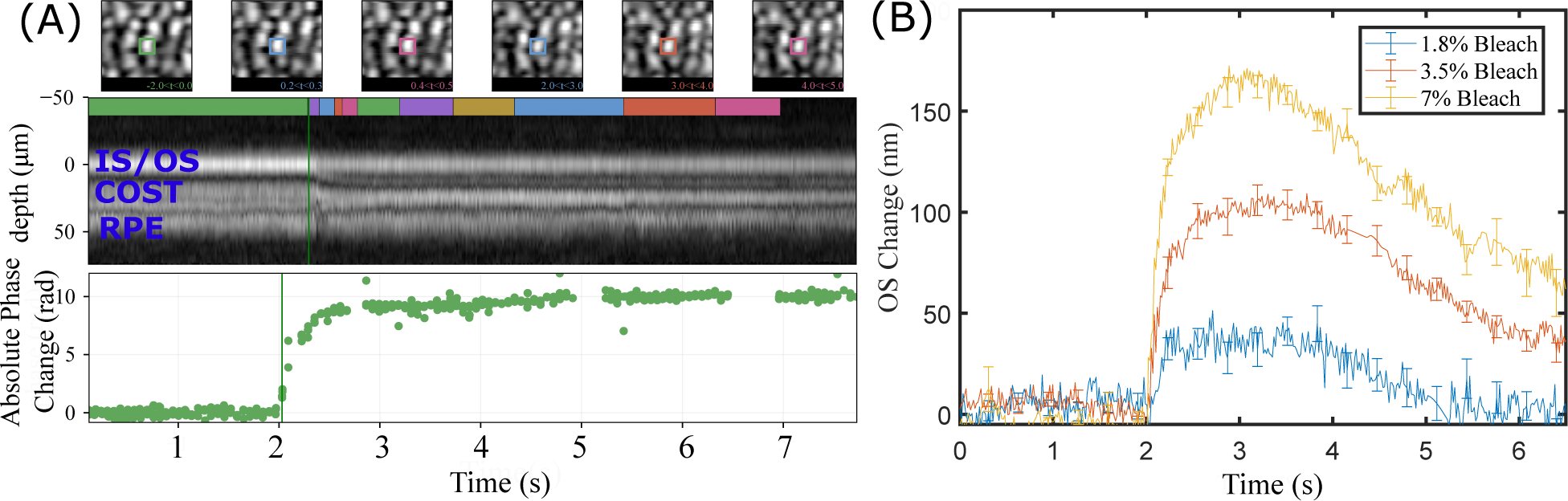
**(A)** Response of a single cone to 70% photopigment bleaching stimuli. The top row shows examples of motion-corrected *en face* projections of the cone’s neighborhood. A time-series of the cone’s axial profile (M-scan) is shown below the *en face* projections, with a green line indicating the stimulus flash. The phase difference between the IS/OS and COST was monitored as a function of time and can be seen in the bottom plot. **(B)** OS length change as a function of time for lower L/M photopigment bleaching percentages of 1.8, 3.5 and 7. Each curve was produced by averaging responses of 10 – 30 cones. Error bars indicate ± one standard deviation.

**Figure 5:**
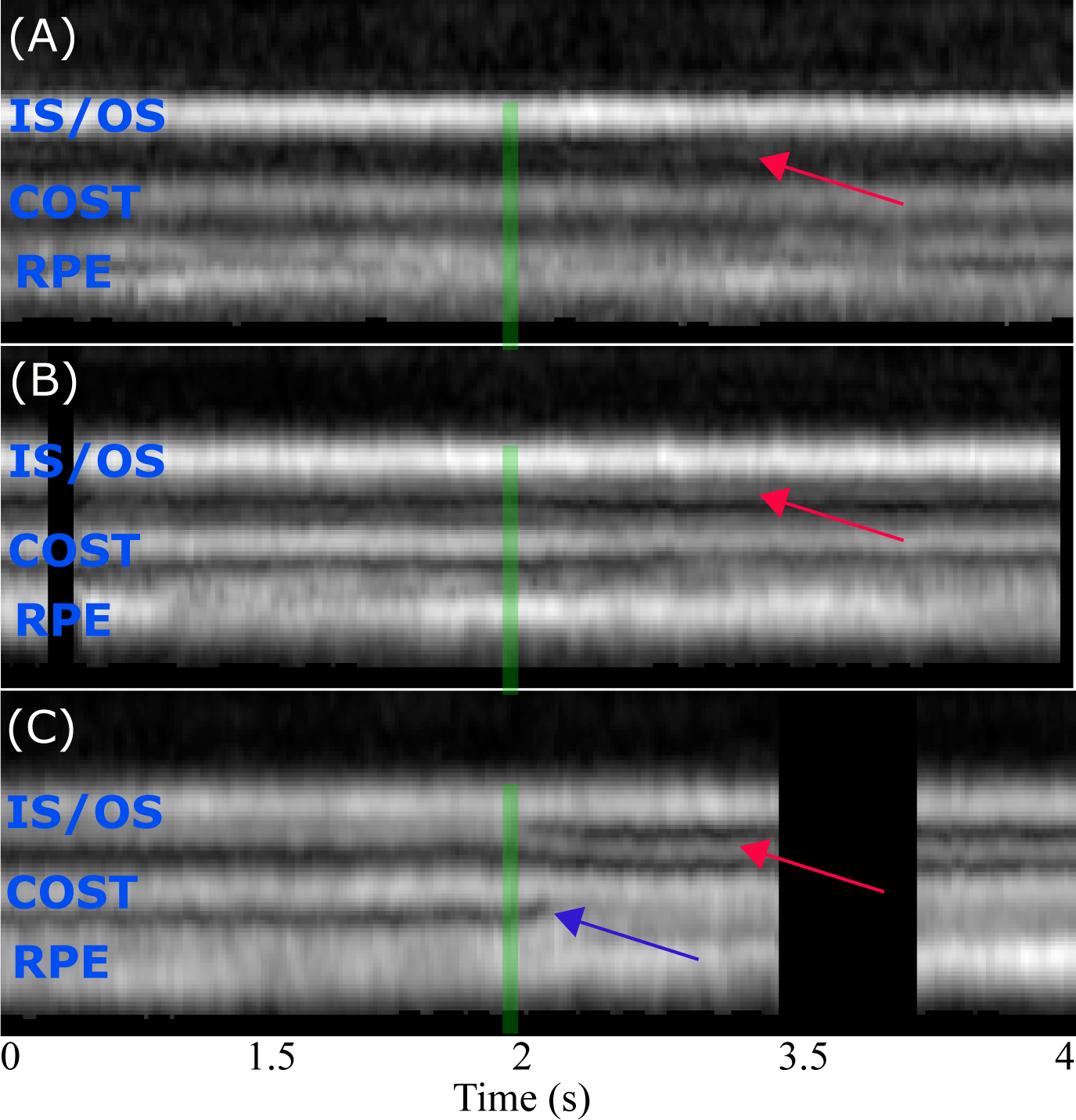
Changes in the axial morphology of cones for photopigment bleaching percentages of **(A)** 1.8, **(B)** 7 and **(C)** 70. As shown by red arrows, appearance of an extra band between IS/OS and COST was observed in most of the cones. The reflectivity of this extra band and also its axial distance from IS/OS seems to be proportional to the bleaching light intensity. The blue arrow indicates changes observed in the RPE and subretinal space.

A critical feature of this system’s design is its speed. For the most intense stimuli, we observed initial phase changes of up to 50 rad/s. In order to correctly unwrap phase, the phase change between consecutive samples should be less than *π* radians. This suggests that the minimal rate at which cones must be imaged, i.e., the minimal volume rate, is 16*Hz*. In the presence of noise, it is likely that a higher rate is required. Our volume rate of 32*Hz* proved sufficient for measuring these fast changes. Another benefit of high-speed acquisition is the reduction of intraframe eye movement artifacts, which permits better registration of frames and tracking of cones.

## 4 Conclusions

We have demonstrated that AO-OCT may be used to detect and measure functional responses of foveal cone photoreceptors. It reveals stimulus-evoked OS phase changes, consistent with OS elongation, similar to those previously detected in foveal cones using AO flood illumination and peripheral cones using full-field SS-OCT, and consistent with elongation observed in human and mouse rods. In addition, our images reveal stimulus-evoked changes in the intensity and organization of the outer retinal bands. These functional responses represent a rich set of biomarkers of photoreceptor function.

## Acknowledgments

The authors gratefully acknowledge the assistance of Susan Garcia.

## References

[1] R. S. Jonnal, J. Rha, Y. Zhang, et al., “In vivo functional imaging of human cone photoreceptors,” Opt. Express 15(24), 16141–16160 (2007).

[2] V. Srinivasan, Y. Chen, J. Duker, et al., “In vivo functional imaging of intrinsic scattering changes in the human retina with high-speed ultrahigh resolution OCT,” Opt Express 17(5), 3861–3877 (2009).

[3] M. D. Abràmoff, R. F. Mullins, K. Lee, et al., “Human photoreceptor outer segments shorten during light adaptation,” Invest Ophth Vis Sci 54(5), 3721–3728 (2013).

[4] C. D. Lu, B. Lee, J. Schottenhamml, et al., “Photoreceptor layer thickness changes during dark adaptation observed with ultrahigh-resolution optical coherence tomography,” Investigative Ophthalmology & Visual Science 58(11), 4632–4643 (2017).

[5] D. Hillmann, H. Spahr, C. Pfäffle, et al., “In vivo optical imaging of physiological responses to photostimulation in human photoreceptors,” Proc. Natl. Acad. Sci. U.S.A. 113(46), 13138–13143 (2016).

[6] Y. Zhang, J. Rha, R. S. Jonnal, et al., “Adaptive optics parallel spectral domain optical coherence tomography for imaging the living retina,” Opt Express 13(12), 4792–4811 (2005).

[7] R. Zawadzki, S. Jones, S. Olivier, et al., “Adaptive-optics optical coherence tomography for high-resolution and high-speed 3d retinal in vivo imaging,” Opt Express 13(21), 8532–8546 (2005).

[8] R. Huber, M. Wojtkowski,, et al., “Fourier domain mode locking (fdml): A new laser operating regime and applications for optical coherence tomography,” Opt. Express 14(8), 3225–3237 (2006).

[9] T. Klein, W. Wieser, C. M. Eigenwillig, et al., “Megahertz oct for ultrawide-field retinal imaging with a 1050 nm fourier domain mode-locked laser,” Opt. Express 9(4), 3044–3062 (2011).

[10] Y. Chen, D. M. de Bruin, C. Kerbage, et al., “Spectrally balanced detection for optical frequency domain imaging,” Opt. Express 15(25), 16390–16399 (2007).

[11] S.-H. Lee, J. S. Werner, and R. J. Zawadzki, “Improved visualization of outer retinal morphology with aberration canceling reflective optical design for adaptive optics-optical coherence tomography,” Biomed Opt Express 4(11), 2508–2517 (2013).

[12] R. S. Jonnal, O. P. Kocaoglu, R. J. Zawadzki, et al., “The cellular origins of the outer retinal bands in optical coherence tomography images,” Invest Ophth Vis Sci 55(12), 7904–7918 (2014).

[13] C. Curcio, K. Sloan, R. Kalina, et al., “Human photoreceptor topography.,” J Comp Neurol 292(4), 497–523 (1990).

[14] S. Stevenson and A. Roorda, “Correcting for miniature eye movements in high-resolution scanning laser ophthalmoscopy,” Proc. SPIE 5688, 145–151 (2005).

[15] R. S. Jonnal, O. P. Kocaoglu, Q. Wang, et al., “Phase-sensitive imaging of the outer retina using optical coherence tomography and adaptive optics,” Biomed Opt Express 3(1), 104–124 (2012).

[16] M. A. Choma, A. K. Ellerbee, C. Yang, et al., “Spectral-domain phase microscopy,” Opt Lett 30(10), 1162–1164 (2005).

[17] P. Zhang, R. J. Zawadzki, M. Goswami, et al., “In vivo optophysiology reveals that g-protein activation triggers osmotic swelling and increased light scattering of rod photoreceptors,” Proc Nat Acad Sci USA 114(14), 2937–2946 (2017).

[18] E. Götzinger, M. Pircher, W. Geitzenauer, et al., “Retinal pigment epithelium segmentation by polarization sensitive optical coherence tomography,” Opt Express 16(21), 16410–16422 (2008).

[19] Q.-X. Zhang, R.-W. Lu, J. D. Messinger, et al., “In vivo optical coherence tomography of light-driven melanosome translocation in retinal pigment epithelium,” Scientific Reports 3, 2644 (2013).

